# Synergistic CDK control pathways maintain cell size homeostasis

**DOI:** 10.1101/2020.11.25.397943

**Authors:** James O. Patterson, Souradeep Basu, Paul Rees, Paul Nurse

## Abstract

To coordinate cell size with cell division, cell size must be computed by the cyclin-CDK control network to trigger division appropriately. Here we dissect determinants of cyclin-CDK activity using a novel high-throughput single-cell in vivo system. We show that inhibitory phosphorylation of CDK encodes cell size information and works synergistically with PP2A to prevent division in smaller cells. However, even in the absence of all canonical regulators of cyclin-CDK, small cells with high cyclin-CDK levels are restricted from dividing. We find that diploid cells of equivalent size to haploid cells exhibit lower CDK activity in response to equal cyclin-CDK enzyme concentrations, suggesting that CDK activity is reduced by DNA concentration. Thus, multiple pathways directly regulate cyclin-CDK activity to maintain robust cell size homeostasis.

## Main

Cells display homeostatic behavior in maintaining population cell size by controlling cell size at cell division. This homeostasis is thought to be driven by ensuring that larger cells are more likely to divide than smaller cells, resulting in the correction of any cell size deviances at cell division^1^. Cyclin dependent kinase (CDK^Cdc2^) is the master regulator of the eukaryotic cell cycle, and therefore the propensity for smaller cells not to divide must feed into the regulation of cyclin-CDK^2^. CDK is subject to several mechanisms of control: cyclin synthesis, and subsequent binding to CDK drives CDK into a catalytically competent form^3^; Wee1 kinase and Cdc25 phosphatase act to inhibit or activate CDK respectively through regulatory tyrosine phosphorylation^4,5^; and PP2A phosphatase works to remove phosphates deposited by CDK, effectively reducing its activity^6^.

Much of the data about CDK regulation has been acquired *in vitro*^7–11^, and the quantitative influence of the known regulatory mechanisms *in vivo* has been less studied. Thus, it remains unclear how cell size information feeds into this regulatory network to prevent smaller cells from division, and thus maintain size homeostasis.

Given the complexity of the CDK regulatory network, we used fission yeast cells containing a reduced CDK control system with the cell cycle being driven by a monomeric cyclin-CDK fusion-protein (C-CDK)^2^. This simplifies the network by eliminating cyclin binding to CDK as a regulatory component, and by allowing co-expression of both cyclin and CDK from a single promoter. Using this system, inhibitory Wee1-dependent phosphoregulation can also be removed using a non-phosphorylatable C-CDK^AF^ mutant. These C-CDK^AF^ strains are viable, co-ordinate cell division with cell growth, and maintain cell-size homeostasis (Fig. 1a)^12^.

**Figure 1:**
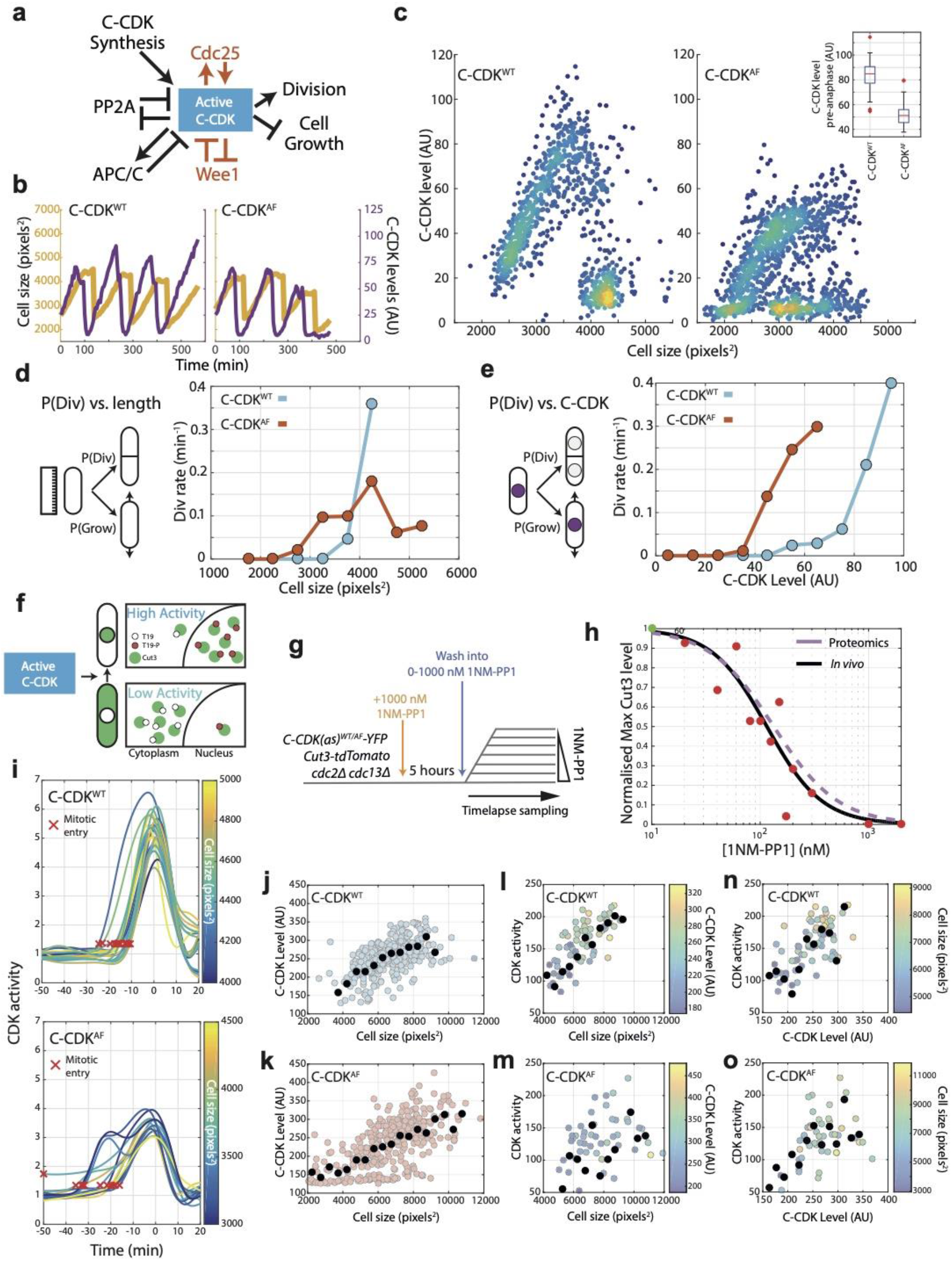
Cell size and C-CDK concentration dictate probability of division and CDK activity in C-CDK^WT^ and C-CDK^AF^ cells. **a** Schematic of major components influencing C-CDK activity at mitosis, and in red the pathways that do not influence C-CDK^AF^. **b** Example cell lineage traces from timelapse microscopy. Cell size in pixels^2^ is given in orange, and C-CDK fluorescence intensity is given in purple. Steep decreases in cell size traces correspond to cell division. **c** Scatter plot of mean C-CDK level vs. cell size from timelapse microscopy data. C-CDK level is a measure of C-CDK fluorescence intensity. Colours indicate density of data. Inset boxplot is mean nuclear C-CDK concentration immediately prior to degradation at anaphase. Boxes represent IQR, with whiskers delimiting 5^th^ to 95^th^ percentiles. C-CDK^WT^ n=28, C-CDK^AF^ n=44 full cycles. **d** Plot of the probability of division at the next timepoint (P(Div)) vs cell length for CDK^WT^ and CDK^AF^. Cells were followed through timelapse microscopy with measurements taken each frame. P(Div) defined as the proportion of cells that undergo C-CDK degradation at anaphase by the next timepoint, given as rate per minute. Points represent cells binned by size, with points plotted at bin centre. C-CDK^WT^ n=685, C-CDK^AF^ n=961 timepoints. **e** Plot of P(Div) function vs C-CDK level for CDK^WT^ and CDK^AF^. C-CDK^WT^ n=685, C-CDK^AF^ n=961 timepoints. C-CDK intensity measurements taken every frame from timelapse microscopy, and binned by C-CDK level. **f** Schematic of Cut3 as a CDK activity reporter. Mitotic CDK dependent phosphorylation of Cut3 on T19 results in nuclear translocation of the protein. **g** Experimental outline of block and release timelapse experiment for panels (h),(j)-(o). Asynchronous cells possessing an analogue sensitive (as) CDK were blocked in G2 using 1 μM 1NM-PP1 for 5 hours, and then released into a range of 1NM-PP1 concentrations. Cells were then followed and monitored for their Cut3-tdTomato nuclear/cytoplasmic (N/C) ratio (C-CDK activity) and C-CDK-YFP level using fluorescence timelapse microscopy (see methods). **h** Maximum CDK activity (normalized against maximum level, obtained by release into DMSO) against 1NM-PP1 concentration. Red points are the median of the data sets for each drug concentration (N=324), green point is median in DMSO. Black line is the Hill equation fit to the median data by a nonlinear fitting algorithm (IC50=115.4, Hill coefficient=-1.71). Purple dashed line is Hill curve derived from Swaffer *et al*. (2016) dose response data (IC50=133.4, Hill coefficient=-1.47). **i** Timelapse quantification of CDK activity in asynchronous cells. Traces are aligned so that 0 minutes corresponds to peak Cut3-tdTomato N/C ratio. Curve smoothing could move Cut3 peak earlier/later than exactly 0 min. Trace colour indicates cell size. Red X indicates automatically defined mitotic entry point. C-CDK^WT^ n=23 and C-CDK^AF^ n=14. **j** Scatter plot of C-CDK-YFP levels against cell size. Experiment described in (g), with measurements taken before release from 1NM-PP1 block. Black points indicate binned data, bin window size 500 pixels^2^. n=324. **k** As in (j), but with C-CDK^AF^, n=312. **l** Scatter plot of peak Cut3 level vs cell size. Experiment described in (g), with measurements taken after release from 1NM-PP1 block into DMSO. Black points indicate binned data, bin window size 500 pixels^2^. Points are coloured by YFP C-CDK levels at release. n=83. R^2^ = 0.5040. **m** As in (l), but with C-CDK^AF^, n=81. R^2^ = 0.2150. **n** Scatter plot of peak Cut3 level vs. C-CDK level after release from 1NM-PP1 block into DMSO. Black points indicate binned data, bin window size 15 AU. Points are coloured by cell size at release. n=83. R^2^ = 0.3668. **o** As in (n), but with C-CDK^AF^, n=81. R^2^ = 0.5501.

To examine the relationship between cell size, C-CDK concentration, and mitosis, we performed quantitative fluorescence time-lapse microscopy on strains expressing fluorescently tagged C-CDK^WT^ and C-CDK^AF^ (Fig 1a-e, Fig. S1a). This analysis showed robust oscillations of C-CDK^WT^ and C-CDK^AF^, with degradation of C-CDK occurring just before cell division (Fig. 1b). C-CDK^AF^ oscillations were more variable, and 5% of C-CDK^AF^ cells trigger C-CDK degradation in the absence of division (Fig. S1), similar to what has been observed in CDK1^AF^ expressing human cells^13^. In both backgrounds, C-CDK concentration scaled with cell size, with C-CDK^WT^ exhibiting a higher amount of C-CDK to enter mitosis compared to C-CDK^AF^ (Fig. 1c). On investigating the links between the probability of a given cell to divide, cell size, and C-CDK level, we found that for C-CDK^WT^ both cell size and C-CDK level reach sharp thresholds at which cell division rates increase (Fig. 1d,e). In the absence of tyrosine phosphorylation, a sharp threshold for C-CDK^AF^ level still exists (Fig. 1e), but is at a lower level than C-CDK^WT^. C-CDK^AF^ cells fail to generate a sharp threshold for cell size, but even without a clear size threshold C-CDK^AF^ cells still restrict smaller cells from division (Fig. 1d).

C-CDK level is not a direct measure of C-CDK activity because of the multiple regulatory networks affecting CDK^8^. To investigate CDK activity, cell size, and C-CDK level at the same time we developed an *in vivo* single-cell assay of CDK activity. We used Cut3, the Smc4 homolog, as a CDK activity biosensor, as it translocates from the cytoplasm into the nucleus upon CDK-dependent phosphorylation of a single site in its N-terminus (Fig. 1f)^14^. Thus, the Cut3 nuclear/cytoplasmic (N/C) ratio can be used to assess CDK activity, a method that has been applied to other protein kinases^15,16^. As a test of this assay, we blocked cells expressing fluorescently tagged Cut3 in the background of a bulky ATP-analogue sensitive C-CDK^2^ using 1NM-PP1, and tracked single cells following their release from G2 arrest into a range of 1NM-PP1 doses (Fig. 1g, Fig. S2). The response of the maximum nuclear Cut3 concentration to 1NM-PP1 was similar to the one measured in our previous phosphoproteomics study^17^, confirming that our sensor reflects *in vivo* CDK activity (Fig. 1h). Given that our sensor reads *in vivo* CDK activity, we examined CDK activity in unperturbed cells. CDK activity, as measured by the Cut3 N/C ratio, rises to a higher level in C-CDK^WT^ cells in comparison to C-CDK^AF^ cells, and progress through mitosis in C-CDK^AF^ cells is slower and more variable (Fig. 1i, Fig. S3).

We next investigated the links between C-CDK protein levels, CDK activity, and cell size in C-CDK^WT^ and C-CDK^AF^ cells, beyond their physiological cell lengths. During the G2/M block (Fig. 1g), cell size and C-CDK enzyme concentration scaled with each other in both backgrounds (Fig. 1j,k). After the release from CDK inhibition, C-CDK^WT^ activity correlated well with both cell size and C-CDK protein level (Fig. 1l,n). However, peak C-CDK^AF^ activity correlated better with protein level than with cell size (Fig. 1m,o). When conducting this experiment using a high throughput assay (Fig. S4, Fig. S5) we observed similar behavior, but this approach clearly illustrated that peak CDK activity in both C-CDK^AF^ and C-CDK^WT^ was heavily size dependent (Fig. S4e). Therefore, CDK tyrosine phosphorylation helps to inform the cell division machinery of size (Fig. 1d,l). However, in the absence of tyrosine phosphorylation, C-CDK^AF^ cells are still able to generate a threshold C-CDK level for division and show size-dependent CDK activity scaling (Fig. 1e,m,o). Thus, they are still able to restrict small cells from dividing.

A complication of the above assay is that cell size scales with C-CDK level^2,18,19^ (Fig. 1c, j, k). To uncouple cell size from C-CDK level, and study if small cells are prevented from entering mitosis due to low C-CDK level or for some other reason, we developed a more flexible single cell CDK assay system. This assay was also based on Cut3 translocation into the nucleus (Fig. 2a) but uses a brighter synthetic C-CDK activity sensor, synCut3-mCherry to allow its co-detection with C-CDK in a high-throughput assay (Fig. S6). This sensor was expressed in a strain where the endogenous CDK network can be switched off using a temperature sensitive CDK1 allele, *cdc2*^*TS*^. A tetracycline-inducible fluorescently-tagged C-CDK was constructed which was made non-degradable^20^ and sensitive to inhibition by 1NM-PP1. Induction of C-CDK at the *cdc2*^*TS*^ restrictive temperature allows the study of the activity of the inducible C-CDK without either wild-type CDK activity or C-CDK proteolysis during. Using this assay, we acquired hundreds of thousands of images of single cells, which allowed us to study the *in vivo* biochemistry of CDK activity in response to a wide range of C-CDK concentrations, in physiologically-sized cells. C-CDK level was uncoupled from cell size as induction of C-CDK was not dependent on cell size (Fig. 2b,c). Results from this assay demonstrated that *in vivo* CDK activity was dependent on C-CDK level, and was reduced when CDK activity was inhibited using 1NM-PP1 (Fig. 2d) (Fig. S7).

**Figure 2:**
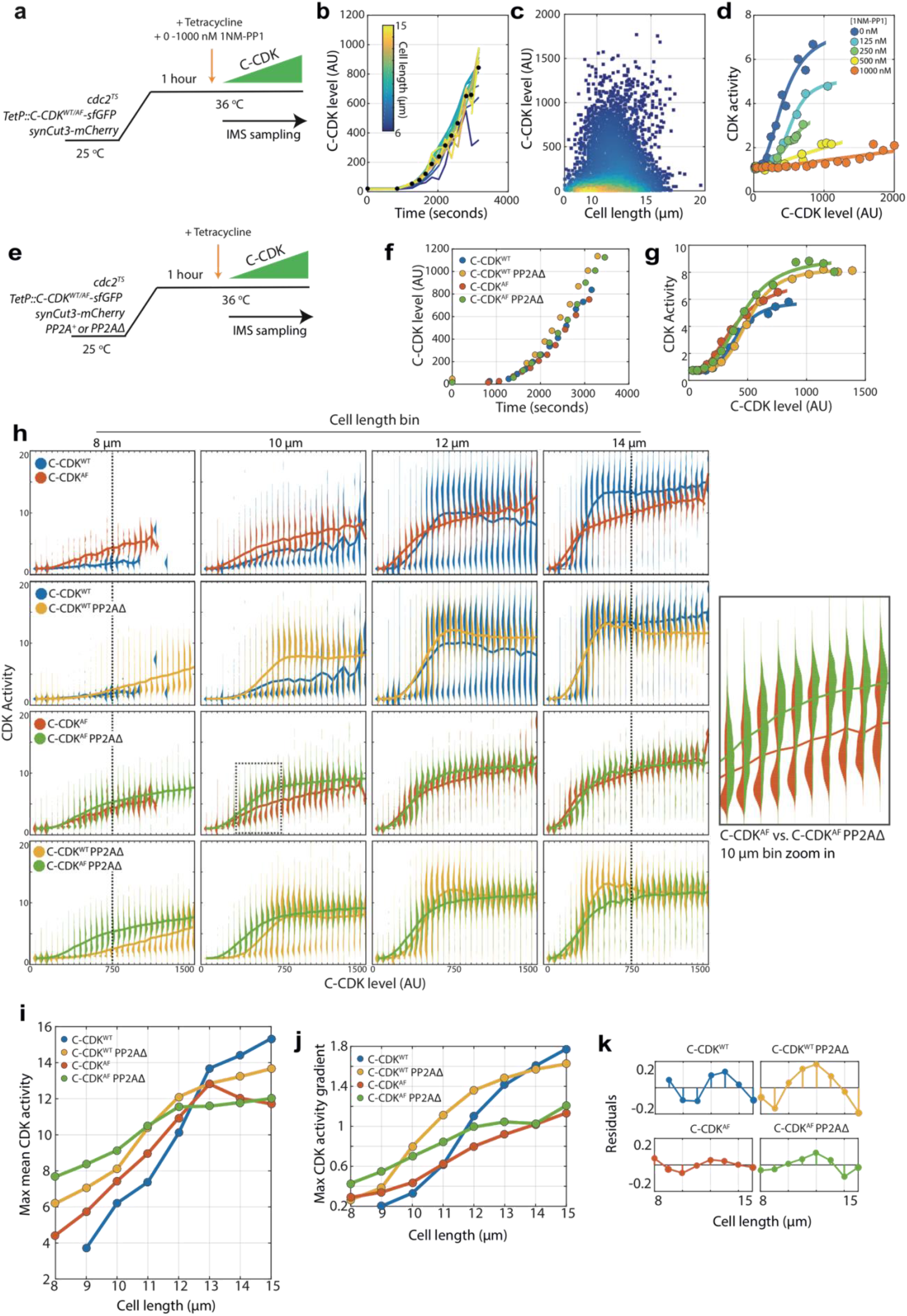
Cell size is able to modulate CDK activity independently of canonical CDK regulation. **a** Experimental outline for figure for panels (b)-(d). Cells were held at 36°C for 1 hour to ablate the function of the temperature sensitive (TS) *cdc2* allele. C-CDK-sfGFP expression was induced by addition of tetracycline, and ectopic C-CDK concentration and CDK activity were measured by sequential sampling during induction. Induced C-CDK lacks its degron box sequence, and therefore is not degraded at anaphase. Sequential sampling during C-CDK induction begins at the point of tetracycline addition, with roughly one sample taken every 3 minutes after the start of C-CDK production. Sampling is conducted using an imaging flow cytometer (IMS). **b** Expression of C-CDK^WT^ from point of tetracycline addition. Different coloured lines represent different size bins. Black dots represent mean C-CDK level over all size bins for given timepoint. After lag period of ∼1000 seconds after tetracycline addition, samples are taken roughly every 3 minutes. n=759633. **c** Scatter plot of cell length vs. C-CDK levels. Coloured by density of data points. Data collected throughout induction. n=759633. **d** Mean CDK activity dose response against C-CDK in the presence of annotated levels 1NM-PP1. Circles represent average CDK activities across all cells from a single sample taken after induction. 0 nM n=166081, 125 nM n=60759, 250 nM n=165128, 500 nM n=135670 and 1000 nM n=231995. **e** Experimental outline for panels (f)-(k). Cells were held at 36°C for 1 hour to ablate *cdc2*^*TS*^ function. After 1 hour, C-CDK^WT^ or C-CDK^AF^ was induced with tetracycline in cells with either PP2A deleted or present. Induced C-CDK lacks its degron box sequence, and therefore is not degraded at anaphase. Sequential sampling during C-CDK induction begins at the point of tetracycline addition, with timepoints taken roughly every 3 minutes after 1000 second lag period in C-CDK induction. **f** Induction of C-CDK after tetracycline addition. Points represent mean concentration of C-CDK across all size bins at indicated time points. CDK^WT^ n=166081. C-CDK^WT^ PP2AΔ n=175247. C-CDK^AF^ n=177292. C-CDK^AF^ PP2AΔ n=174847. **g** C-CDK activity against C-CDK level in given genetic backgrounds defined in (f). Points represent mean C-CDK activity of all cells. Data is pooled from experiment in (e), from all time points following tetracycline induction. Key is the same as (f). **h** Violin plots of single cell C-CDK level against CDK activity in annotated size bins and strain backgrounds. Solid line through violin plot indicates the mean CDK activity within the C-CDK level bin. **i** Maximum mean CDK activity vs. cell length in annotated strain backgrounds. Max mean CDK activity is the maximum mean CDK activity within a C-CDK fluorescence level bin for a given cell size. The mean CDK activity level across all fluorescence bins is shown by the solid line in the violin plots in panel (h). **j** Maximum gradient of the mean lines in panel (h) plotted against cell length. Maximum gradient of change is derived from a spline fit to the mean CDK activity vs. C-CDK level trace. **k** Linear regression lines were fit to data in (j), and residuals were plotted (actual value – predicted value). Non-linear residuals indicate bistability in CDK activation.

Combining this system it with genetic backgrounds in which canonical C-CDK regulation was absent, we analysed how mechanisms of CDK regulation affected C-CDK activity in relation to cell size. We performed the assay in backgrounds lacking PP2A, inhibitory CDK tyrosine phosphorylation, or both (Fig. 2e). C-CDK levels increased similarly upon induction in all mutant backgrounds (Fig. 2f). Population mean C-CDK activity was comparable between all conditions (Fig. 2g), however displayed striking differences at the single-cell level when CDK activity was measured in cells of different sizes. In all genetic backgrounds, at the same level of C-CDK enzyme, maximum C-CDK activity increases with cell size (Fig. 2h). This is particularly noticeable when directly comparing the maximum C-CDK activity of cells with C-CDK level of ∼750 AU in the 8 μm bin to the 14 μm bin in all backgrounds (Fig. 2h, dashed lines). The single cell dose-response of CDK activity on C-CDK concentration in a wild-type background is clearly bistable, with cells existing in either an ‘on’ or an ‘off’ state. The C-CDK concentration required to switch cells “on” decreases with increasing cell size, and the sharpness of the transition increases with size (Fig. 2h,j). This bistable behavior is heavily dependent on CDK tyrosine phosphorylation (Fig. 2h,j,k). Removal of PP2A allows the attainment of the “on” state at lower cell sizes (Fig. 2h), effectively shifting the C-CDK dose response curve towards lower sizes without altering the shape of the response (Fig. 2j). In addition, PP2A also adds switch like behavior to the C-CDK activity dose-response, as bistable behavior present with C-CDK^AF^ is not present with C-CDK^AF^ PP2AΔ (Fig. 2h dashed box, inset and 2k).

When looking across all size bins, maximum C-CDK activity increases with cell size in all genetic backgrounds, but plateaus at about 12-13 μm in the absence of tyrosine phosphorylation (Fig. 2i). However, it is clear that cell size is able to regulate C-CDK activity even in the absence of both tyrosine phosphorylation and PP2A (Fig. 2h,i). These results are consistent with our previous observations (Fig. 1), that although tyrosine phosphorylation has a role in informing the cell cycle machinery of size, small cells are still restricted from mitosis even in the absence of tyrosine phosphorylation.

PP2A and inhibitory tyrosine phosphorylation constitute two fundamentally different modes of lowering CDK activity, however it is unknown if they act independently or synergistically to do so. We therefore sought to calculate the individual contributions of PP2A and tyrosine phosphorylation in restricting CDK activity in order to examine if their combined contribution was greater than the sum of their parts. To calculate the individual contributions of tyrosine phosphorylation and PP2A in restricting C-CDK activity, first we measured the threshold C-CDK level required for 50% of cells to reach a C-CDK activity >5 in different strain backgrounds within different size bins (Fig. 3a). This value was chosen as an approximate value of the C-CDK concentration required *in vivo* to trigger mitotic entry in wild-type cells (Fig. 1i). When this C-CDK threshold level was plotted across all size bins (Fig. 3b) the threshold was seen to be size dependent in all strain backgrounds, with wild-type cells exhibiting the strongest capacity to raise the C-CDK level threshold for mitosis in smaller cells. By subtracting the curves of cell length vs. mitotic C-CDK level (Fig. 3c) for various backgrounds we were able to estimate the individual contribution of tyrosine phosphorylation and PP2A in a given background. For example, C-CDK^WT^ PP2AΔ – C-CDK^AF^ PP2AΔ, estimates the ability of tyrosine phosphorylation alone to restrict mitotic entry in a background lacking PP2A. PP2A is able to restrict cells with 600 units of C-CDK from entering mitosis at 8 μm cell length, but only 200 units of C-CDK at 10 μm (Fig. 3c, yellow). If the different components of the CDK control network act separately, adding individual threshold contributions together would generate a threshold curve similar to the wild-type curve. However, when the individual contributions of tyrosine phosphorylation and PP2A, were added to the C-CDK^AF^ PP2AΔ curve, they did not recapitulate the wild-type curve (Fig. 3d). Thus, this analysis demonstrates that there is synergy between the tyrosine phosphorylation network and PP2A activity, and that this synergy is important for establishing the C-CDK level threshold for division.

**Figure 3:**
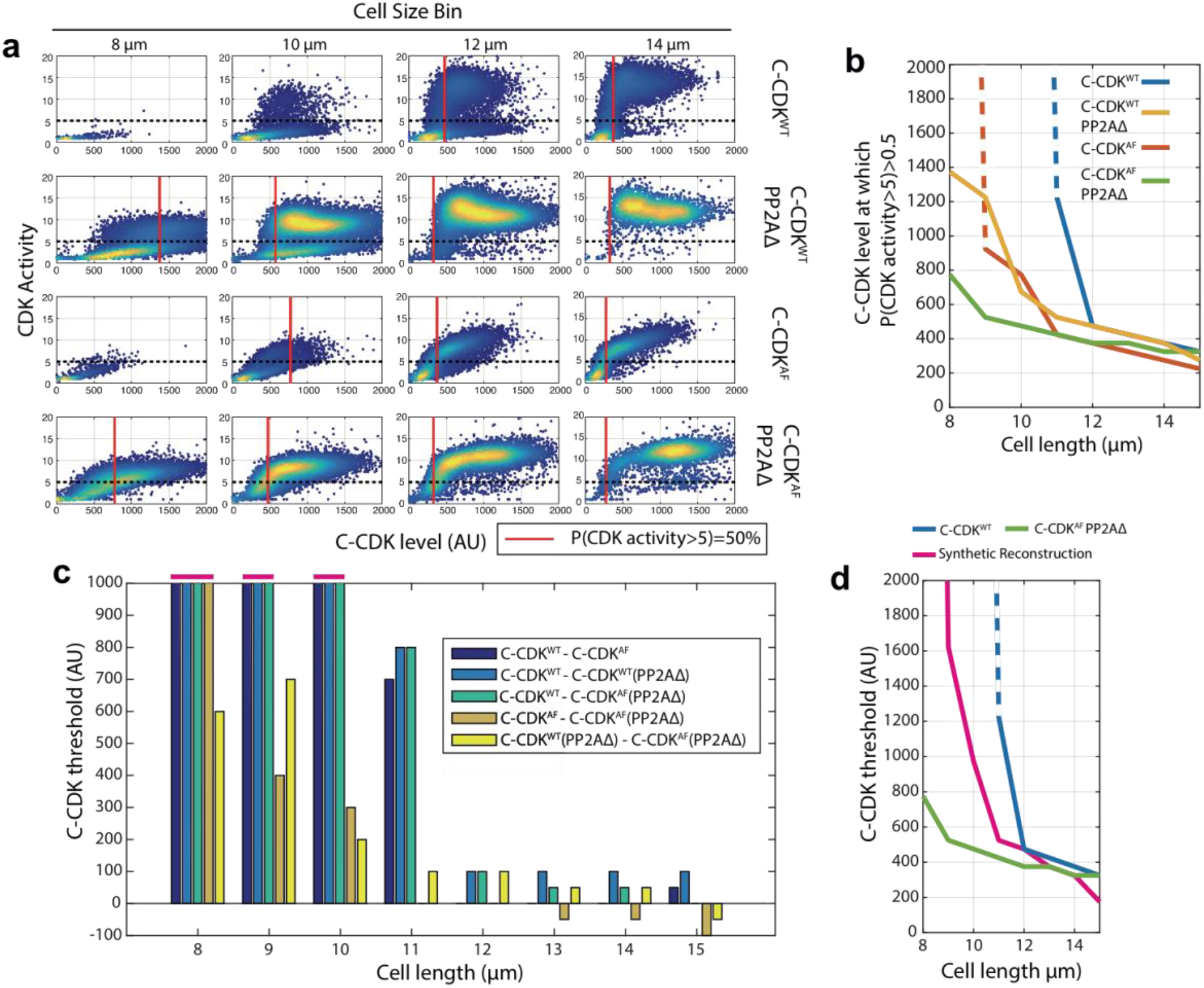
CDK Tyrosine phosphorylation and PP2A act synergistically to restrict division in small cells. **a** Scatter plots of C-CDK level against CDK activity. Either C-CDK^WT^ or C-CDK^AF^ was induced in backgrounds with PP2A either lacking or present. Red line indicates the C-CDK level at which 50% of cells have a CDK activity greater than 5. Black dashed line marks CDK activity of 5. Data taken from Fig. 2h. **b** C-CDK level at which 50% of cells have C-CDK activity > 5. Data is taken from (a) across all size bins. Y-axis represents the C-CDK threshold at which 50% of cells will have a C-CDK activity of 5. Dashed lines indicate values where this C-CDK threshold level is undefined due to the threshold being unattainable in experimental conditions. **c** Piecewise dissection of the amount of C-CDK a particular component of the cell cycle network is able to prevent from switching to an ‘on’ state (C-CDK activity level of 5) in different size bins. Bar chart shown is of subtractions of curves described in key (from inset). For example, C-CDK^WT^ - C-CDK^AF^ gives the C-CDK threshold tyrosine phosphorylation alone (in a background with PP2A present) is able to generate to restrict C-CDK activation. Values that are undefined due to undefined original threshold values from (a) are taken to be 1000 units, and are marked above the axis (pink). **d** Cell length against C-CDK level threshold of annotated curves. Here, a synthetic threshold curve is built (pink), by adding the individual component regulatory contributions of CDK tyrosine phosphorylation (panel (c), yellow) and PP2A (panel (c), orange) to the base curve of C-CDK^AF^ PP2AΔ (green) to try and re-capitulate the WT behaviour (blue). Dashed line indicates undefined threshold values.

We have shown that small cells are normally prevented from division by their low C-CDK protein level (Fig. 1) along with PP2A and tyrosine phosphorylation working synergistically to increase the level of C-CDK needed to trigger division in smaller cells (Fig. 3). Strikingly however, in the absence of these canonical regulators, small cells are still able to restrict division by lowering CDK activity as a result of some other factor related to cell size (Fig. 2h,i,j). This unknown factor is able to lower CDK activity in small cells despite high C-CDK levels, thus restricting them from division (Fig 2i).

Given the positive relationship between maximum C-CDK activity and increasing cell size in the C-CDK^AF^ PP2AΔ mutant (Fig. 2i), we hypothesized that a titration based model might be operative, where cells dilute a CDK inhibitor as they grow^21^. Given that cell size is linked to ploidy through an unknown mechanism, we tested whether DNA concentration could influence CDK activity, and therefore constitute the unknown factor able to lower C-CDK activity in small cells. We induced C-CDK^AF^ in haploid and diploid variants of the C-CDK^AF^ PP2AΔ strain, thereby eliminating all major canonical CDK regulation at mitosis (Fig. 4a,b). Strikingly, diploid cells exhibited lower C-CDK activity in response to the same C-CDK enzyme concentration as haploids (Fig. 4c). The EC50 of the diploid dose response curve was almost double that of the haploid (Fig. 4d). Looking at single-cell, volume-resolved data, the inhibition of C-CDK activity is most marked in smaller diploid cells, with larger diploid cells having almost indistinguishable dose-response curves from their haploid equivalents (Fig. 4e). The effect of cell size on CDK activation is much less marked in these larger than normal haploids (Fig. 4f). The diploids, which feature cells of physiological diploid size, still experience DNA concentration dependent inhibition of their CDK activity. The effect of equal C-CDK levels resulting in lower C-CDK activity in small diploids when compared to equivalent haploids is readily seen from raw images (Fig. 4g). Therefore, in search of additional C-CDK regulation we show that cells of different ploidies, but otherwise equivalent volume, experience variable C-CDK activity in response to equal C-CDK level. This suggests that even in the absence of all canonical CDK regulation, DNA itself is able to lower CDK activity to prevent division in small cells. This regulation appears to operate in a titration-based manner, as at higher volumes this inhibition of CDK activity disappears.

**Figure 4:**
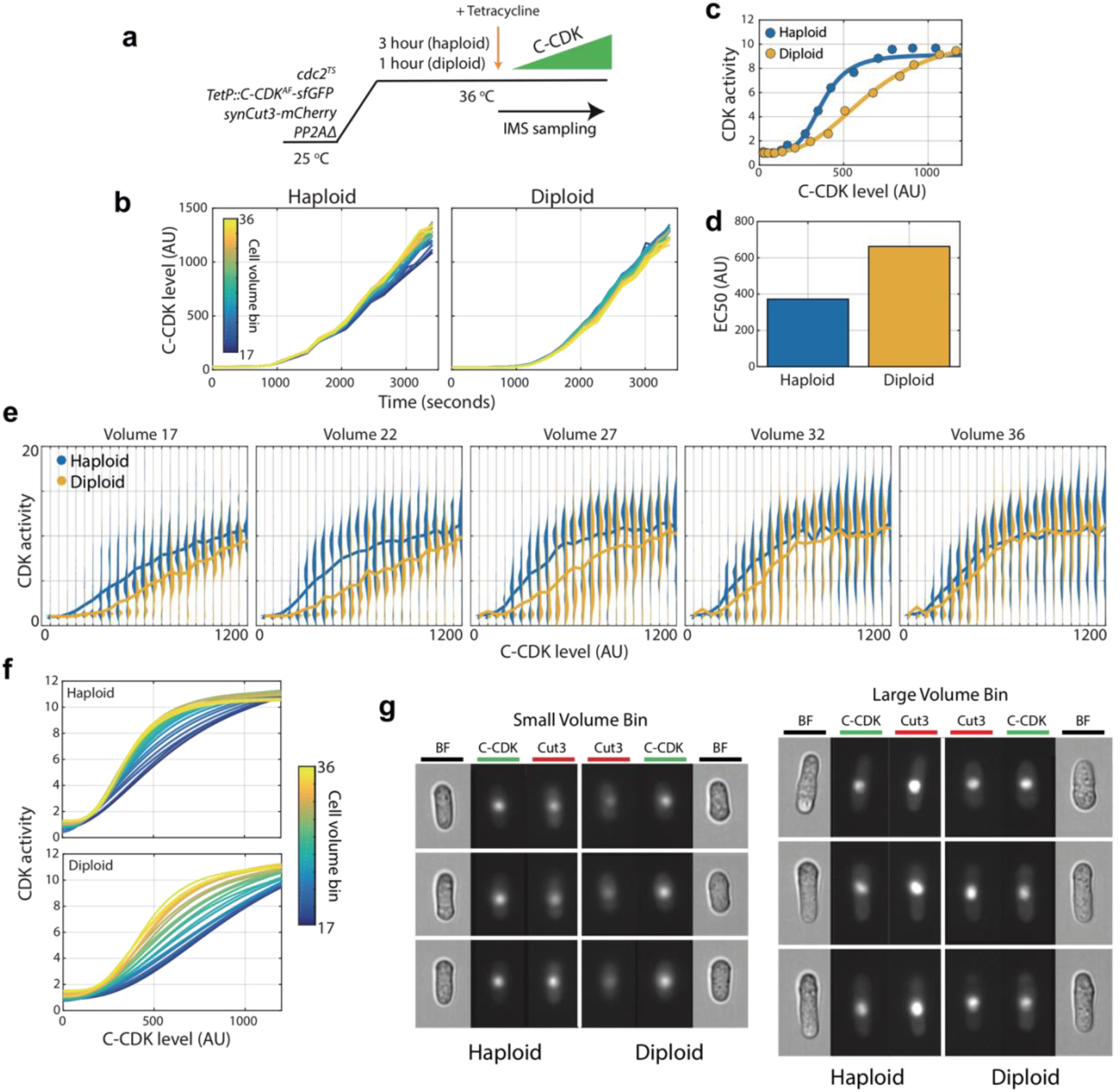
Cellular DNA content inhibits CDK activity independently of tyrosine phosphorylation or PP2A activity. **a** Experimental outline for panels (b)-(g). PP2A Δ/Δ diploids and PP2AΔ haploids were arrested using *cdc2*^*TS*^. Diploids were held at 36°C for 1 hour, whilst haploids were held for 3 hours to generate blocked cell populations with similar cell volumes despite ploidy differences. C-CDK^AF^ expression was induced by addition of tetracycline, and C-CDK concentration and CDK activity were measured by sequential sampling from time of induction in an imaging flow cytometer. **b** Expression of C-CDK^AF^ from point of tetracycline addition in haploid and diploid strains. Different coloured lines represent different size bins. Haploid n=125021, Diploid n=139557. **c** Mean CDK activity against C-CDK^AF^ level in haploids and diploids. Solid line is a sigmoid fit to data. **d** EC50 from sigmoid curves in (c). Haploid EC50: 372 AU. Diploid EC50: 663 AU. Haploid EC50 is 56% of diploid EC50. **e** Violin plots of single cell C-CDK^AF^ level against CDK activity in annotated volume bins and ploidy status. Solid line through violin plot indicates the mean CDK activity within the C-CDK level bin. Volume bins span a physiological range of diploid cell sizes. Volume bin 17 corresponds to a haploid cell length of 12.1 μm and a diploid cell length of 9.53 μm. Volume bin 36 corresponds to a haploid length of 18.7 μm and a diploid length of 14.4 μm. **f** Mean intra volume-bin dose response of C-CDK level vs. CDK activity in annotated ploidy level. Lines are sigmoid curves fit to raw data. Cell volume bin indicated by line colour. **g** Example raw images from experiment. Brightfield (BF) channel displaying cell morphology, C-CDK-sfGFP channel and synCut3-mCherry CDK activity indicator are shown. C-CDK level is the same across all images.

Our approach has demonstrated that three mechanisms contribute to cell size homeostasis through CDK activity control: C-CDK enzyme concentration scaling, synergistic PP2A and tyrosine-phosphorylation dependent C-CDK threshold scaling, and DNA concentration dependent inhibition of C-CDK enzyme activity. Our results demonstrate that C-CDK activity vs. C-CDK level dose-response curves previously demonstrated *in vitro* operate *in vivo*, but in addition we show they are strongly dependent on cell size *in vivo*. We also demonstrate a direct link between ploidy and CDK activity, thus suggesting an explanation for why cell size is linked to ploidy universally across cell types^22–26^. Finally, we show that tyrosine phosphorylation, PP2A activity, and DNA dependent inhibition of CDK activity act together to restrict small cells from division, forming a mechanism to generate the robust cell size threshold behavior observed in normal cells. Cancers often exhibit increased variability in their cell size at division^27^, and further work on which of the three cell size control mechanisms are lost within these tumors could provide a route into developing synthetic lethal approaches by inhibition of the remaining active pathways.

## Methods

### S. pombe genetics and cell culture

S. pombe media and standard methods are as previously described^28^. After nitrogen and glucose addition, EMM was filter sterilised. This process allows for the generation of clear un-caramelised media. Nutritional supplements for auxotrophic yeast strains were added at a concentration of 0.15 mg/ml. Temperature-sensitive mutant strains were grown at temperatures as specified in the text. The temperature-sensitive allele of Cdc2 used was Cdc2-M26. To modulate inducible promoters, anhydrotetracycline (Sigma) in DMSO at specified concentrations was added to 0.03125 μg/ml final concentration unless otherwise specified. To alter Cdc2(as) activity, 1NM-PP1 diluted in DMSO was used at concentrations specified in the text. To stain for septa, calcofluor (Fluorescent Brightener 28 (Sigma Aldrich)) was made up in water at 1 g/L and used as 500x stock. Bortezomib was added to cultures to inhibit the C-CDK degradation, as described previously^29^. SynCut3 was constructed by Gibson assembly of a codon optimised fragment consisting of the first 528 amino acids of Cut3, a linker region, and a fluorescent protein (mCherry or mNeongreen). YFP was tagged onto C-CDK at the C-terminus of the protein. Where the sfGFP labelled C-CDK was used, the sfGFP was present internally within the Cdc13 component^29^. Cut3-mCherry was generated by C-terminal tagging^30^ and Cut3-GFP was developed previously^14^. Details of the TetR promoter and linearised variants can be found in a previous publication^1^.

### Imaging flow cytometry

Imaging flow cytometry was performed using an Imagestream Mark X two-camera system (Amnis), using the 60x objective. Cells were concentrated by centrifugation (5000 rpm/30 seconds) and resuspended in ∼25 μl of media before sonication in a sonicating water bath.

1. Gradient RMS>65 (a measure of cell focus).
2. Area/Aspect ratios consistent with single cells.

To avoid any autofocus based drift within an experiment, cell were imaged at fixed, empirically determined focal points, designed to maximise the number of cells with gradient RMS>65. Data was analysed using custom Matlab scripts.

To perform time-lapse imaging flow cytometry, water baths at specified temperatures for the experiment were set up with cultures next to the IMS. Time was measured from the point of drug addition to liquid culture or as described during a wash protocol for drug release. Samples were collected as above from the waterbath, and sample time-points defined as the time at which acquisition on the IMS began (as opposed to time when sample was collected – although this was consistently ∼3 minutes apart). Samples were imaged for ∼1 minute unless otherwise stated.

### Microscopic imaging

All imaging was performed using a Deltavision Elite (Applied Precision) microscope – an Olympus IX71 wide-field inverted fluorescence microscope with a PLAN APO 60x oil, 1.42 NA objective and a Photometrics CoolSNAP HQ2 camera. To maintain specified temperatures during imaging, an IMSOL imcubator Environment control system and an objective heater was used. SoftWoRx was used to set up experiments. 5 z-stacks were acquired, with 1 μm spacing. Image analysis was performed using custom Matlab scripts.

The ONIX Microfluidics platform allows for long-term time-lapse imaging of live cells. Plate details can be found at http://www.cellasic.com/ONIX_yeast.html. 50 μl of cell culture at density 1.26×106/ml was loaded into the plate, and imaged in the 3.5 μm chamber. Cells were loaded at 8 psi for 5 seconds. Media was perfused at a flow rate of 3 psi. The imaging chamber was washed with media for 1 minute at 5 psi before cells were loaded.

Mattek glass bottom dishes were used for some time-lapse imaging applications with drugs that were incompatible with Cellasics plates, primarily for the purpose of release from a 1NM-PP1/Cdc2(as) cell cycle block. Dishes were pre-treated with soybean lectin to permit cell adherence (Sigma Aldrich). Before addition of cells Mattek dishes were pre-warmed on a heatblock at appropriate temperature. Cells were grown and blocked in liquid culture before 2 ml were pelleted (5000 rpm/30 seconds). Cell pellets were then pooled and resuspended in 1 ml of release media (at which time a stop watch was started) in a new microcentrifuge tube before pelleting (5000 rpm/30 seconds) and resuspended in 5 μl of media. This concentrated cell suspension was then applied to the centre of the Mattek dish, and allowed to settle for ∼5 seconds. The dish was then washed with 1 ml of release media 3x. The dish was then filled with 3 ml of release media before rapid imaging. In general the wash process requires 1.5 minutes, and imaging setup requires 5 minutes for ∼8 FOV.

## Data analysis and plotting

### Boxplots

The top of box is the 25th percentile of the data, the bottom is the 75th percentile. The line in the middle of the box is the median. Whisker lengths are either the distance to the furthest point outside of the box, or 1.5x the interquartile range, whichever is lower. If data exists that is greater than 1.5x the interquartile range from the top or bottom of the box, this is shown as a red “+”.

### Statistical testing

Statistical testing was performed where appropriate using a two tailed two sample t-test. P values below 0.05 were considered significant. Replicates are shown where appropriate by N numbers.

### Cell size measurement

Cell size was measured by three different metrics. In timelapse microscopy assays, cell size was determined as the area of the 2D surface segmented by our segmentation algorithm. In the high-throughput imagestream assays, cell size was measured as length of the cell. The difference in metric choice between these two systems was due to improved ability of measuring cell length in the high-throughput assay, where it was less affected by focal dependent changes in cell volume. In the haploid vs. diploid experiments, a measure of cell volume was used, where cells were assumed to behave as cylinders, and volume was calculated from the measured radius and length. This was done as diploids are wider than haploids and thus a simple length metric cannot be employed for size binning.

## Strain table

**Strain table:**
| Strain ID | Strain genotype | Source |
| --- | --- | --- |
| JP223 | h? leu1::cdc13P:cdc13-cdc2.as-YFP:cdc13T::ura4 cdc13Δ::natMX6 cdc2::scLeu2 | This work |
| JP224 | h? leu1::cdc13P:cdc13-cdc2AF.as-YFP:cdc13T::ura4 cdc13Δ::natMX6 cdc2::scLeu2 | This work |
| JP670 | h? leu1::cdc13P:cdc13-cdc2.as-YFP:cdc13T::ura4 cdc13Δ::natMX6 cdc2::scLeu2 ura4::Ppcna1-CFP-pcna cut3-mCherry::hphMX6 | This work |
| JP671 | h? leu1::cdc13P:cdc13-cdc2AF.as-YFP:cdc13T::ura4 cdc13Δ::natMX6 cdc2::scLeu2 ura4::Ppcna1-CFP-pcna cut3-mCherry::hphMX6 | This work |
| JP310 | h? leu1::cdc13P:cdc13-cdc2.as:cdc13T::ura4 | This work |
|  | cdc13Δ::natMX6 cdc2::scLeu2<br>cut3-tdTomato::hphMX6 |  |
| JP311 | h? leu1::cdc13P:cdc13-cdc2AF.as:cdc13T::ura4<br>cdc13Δ::natMX6 cdc2::scLeu2<br>cut3-tdTomato::hphMX6 | This work |
| JP295 | h? leu1::cdc13P:cdc13-cdc2AF.as:cdc13T::ura4<br>cdc13Δ::natMX6 cdc2::scLeu2<br>cut3-GFP::ura4 | This work |
| JP296 | h? leu1::cdc13P:cdc13-cdc2.as:cdc13T::ura4<br>cdc13Δ::natMX6 cdc2::scLeu2<br>cut3-GFP::ura4 | This work |
| JP501 | h? cdc2.as::blastMX6 synCut3-<br>mNeongreen:: leu1+ | This work |
| JP507 | h? cdc2.as::blastMX6 synCut3-<br>T19V-mNeongreen:: leu1+ | This work |
| JP601 | h? synCut3-mCherry:: leu1+ cut3-<br>GFP::ura4 | This work |
| JP602 | h? cdc2.as::blastMX6 synCut3-<br>mCherry::leu1+ cut3-GFP::ura4 | This work |
| JP591 | h? cdc2-M26 synCut3-<br>mCherry::leu1+<br>leu1::enoTetP:cdc13-sfGFP-<br>cdc2.as:adh1T::hphMX6 TetR1 | This work |
| JP593 | h? cdc2-M26 synCut3-<br>mCherry::leu1+<br>leu1::enoTetP:cdc13-sfGFP-<br>cdc2AF.as:adh1T::hphMX6 TetR1 | This work |
| JP603 | h? cdc2-M26 synCut3-<br>mCherry::leu1+<br>leu1::enoTetP:DBΔcdc13-sfGFP-<br>cdc2.as:adh1T::hphMX6 TetR1 | This work |
| JP605 | h? cdc2-M26 synCut3-<br>mCherry::leu1+ (JPp178)<br>leu1::enoTetP:DBΔcdc13-sfGFP-<br>cdc2AF.as:adh1T::hphMX6 TetR1 | This work |
| JP679 | h? cdc2-M26::blastMX6 synCut3-<br>mCherry::leu1+<br>leu1::enoTetP:DBΔcdc13-sfGFP-<br>cdc2.as:adh1T::hphMX6 TetR1<br>PP2AΔ::kanMX6 | This work |
| JP680 | h? cdc2-M26::blastMX6 synCut3-<br>mCherry::leu1+<br>leu1::enoTetP:DBΔcdc13-sfGFP-<br>cdc2AF.as:adh1T::hphMX6 TetR1<br>PP2AΔ::kanMX6 | This work |

## Supplementary Figures

**Supplementary Figure 1:**
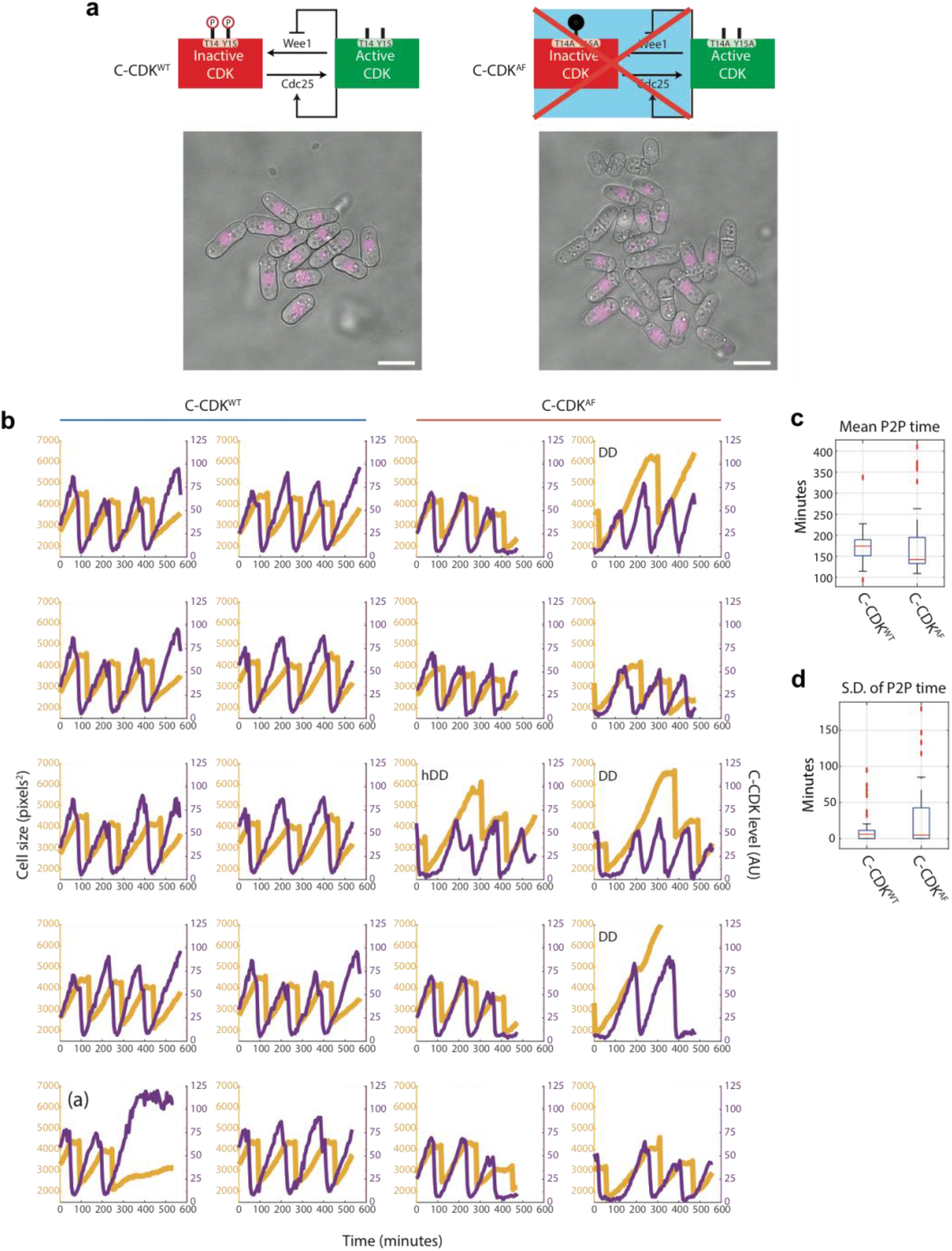
Fluorescence time-lapse quantification of C-CDK dynamics in unperturbed cell cycles. **a** Schematics of C-CDK^WT^ and C-CDK^AF^ regulation by Wee1 kinase and Cdc25 phosphatase. C-CDK^AF^ has T14 mutated to A and Y15 mutated to F to mimic constitutive dephosphorylation of both residues. Example images of a FOV from time-lapse movie is shown. Cells were grown in a Cellasics microfluidics plate following 2 days of culture in YE4S at 32 °C. C-CDK-YFP is seen in purple. Scale bar=10 μm. **b** Purple lines indicate C-CDK levels (mean nuclear concentration) and yellow indicates cell size (measured by cell mask area in pixels^2^). Cell mask and lineage tracing generated by Pomseg and Pomtrack (see methods). DD=Double dip cell, hDD=half double dip cell. DD cells undergo complete cyclin degradation without cell division. hDD cells undergo incomplete cyclin degradation without division. Trace marked (a) represents an abberant cycle in a C-CDK^WT^ expressing cell. **c** Boxplot of C-CDK oscillation period. Period was calculated by measuring the peak to peak (P2P) distance on the autocorrelation function of each C-CDK level lineage trace. C-CDK^WT^, N=32; C-CDK^AF^, N=57. Box represents median value delimited by 25^th^ and 75^th^ percentiles. See methods for outlier points. **d** Boxplot of intra-lineage standard deviation of period length. C-CDK^WT^, N=32; C-CDK^AF^, N=57. Box represents median value delimited by 25^th^ and 75^th^ percentiles. See methods for outlier points.

**Supplementary Figure 2:**
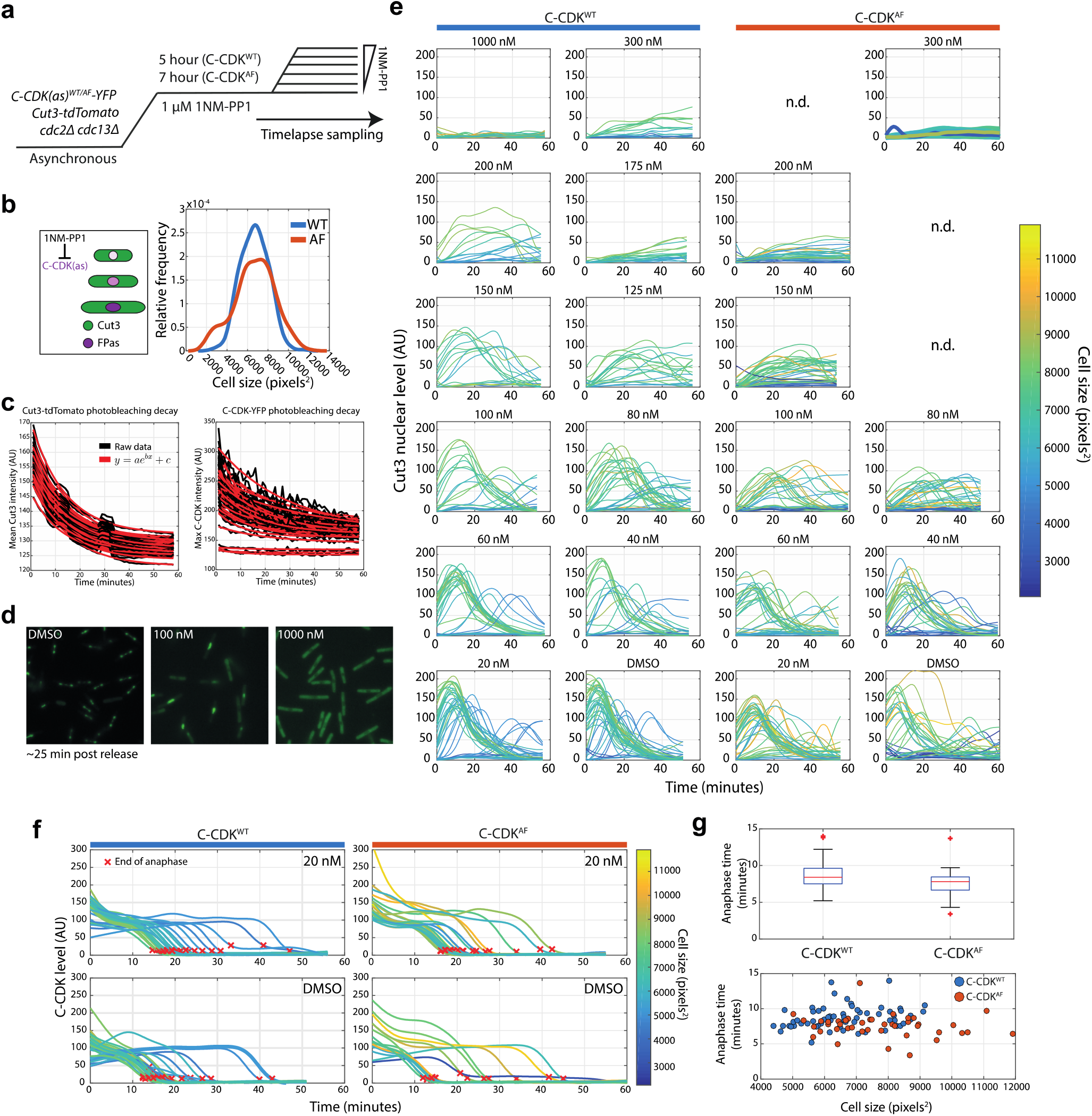
A time-lapse block and release assay to measure the effect of CDK inhibition on CDK activity in single cells. **a** Experimental outline for panels B-G. 1NM-PP1 sensitive C-CDK^WT^ and C-CDK^WT^ cells are blocked by addition of 1NM-PP1. C-CDK^AF^ cells were block for longer (7 hours against 5 hours) to allow cells to reach a similar size distribution as C-CDK^WT^ cells. Cells were then released into a range of 1NM-PP1 concentrations. After release, images were acquired every minute. Time between washing and image acquisition is ∼5 minutes. Cells were grown in EMM at 32°C. **b** Left: Schematic demonstrating that as cells are blocked at G2/M, they continue to grow and accumulate C-CDK but do not translocate Cut3 into the nucleus or alter their levels of Cut3. Right: Density plot demonstrates the overlap population cell lengths of C-CDK^WT^ and C-CDK^WT^ cells after variable block times. **c** Black traces indicate raw data. Red traces indicate exponential curve fit to data. Photobleaching curves were derived from the 1000 nM release using C-CDK^WT^-YFP and Cut3-tdTomato. All subsequent measurements were corrected for photobleaching from derived curves. **d** Images of Cut3-GFP channel from representative FoV ∼25 minutes after release from a 1 μM block into indicated drug concentrations. **e** Plots of nuclear Cut3-GFP levels against time after release over a range of 1NM-PP1 concentrations. Lines are coloured by cell size at T=0 of the release. **f** Single cell C-CDK-YFP traces in DMSO and 20 nM of release. Red x indicates end of anaphase. Traces are coloured by cell size at Time=0. Only traces which undergo anaphase are shown. End of anaphase defined as first time-point at which C-CDK-YFP trace is equal to post anaphase YFP plateau level +10 AU. **g** Boxplot of anaphase time in WT and AF strains. Anaphase time is calculated as end of anaphase time – peak Cut3 time. Difference is non-significant. C-CDK^WT^, N=69 and C-CDK^AF^, N=47. Lower panel, scatter plot of anaphase time vs cell size, with strain indicated by colour. Box represents median value delimited by 25^th^ and 75^th^ percentiles. See methods for outlier points.

**Supplementary Figure 3:**
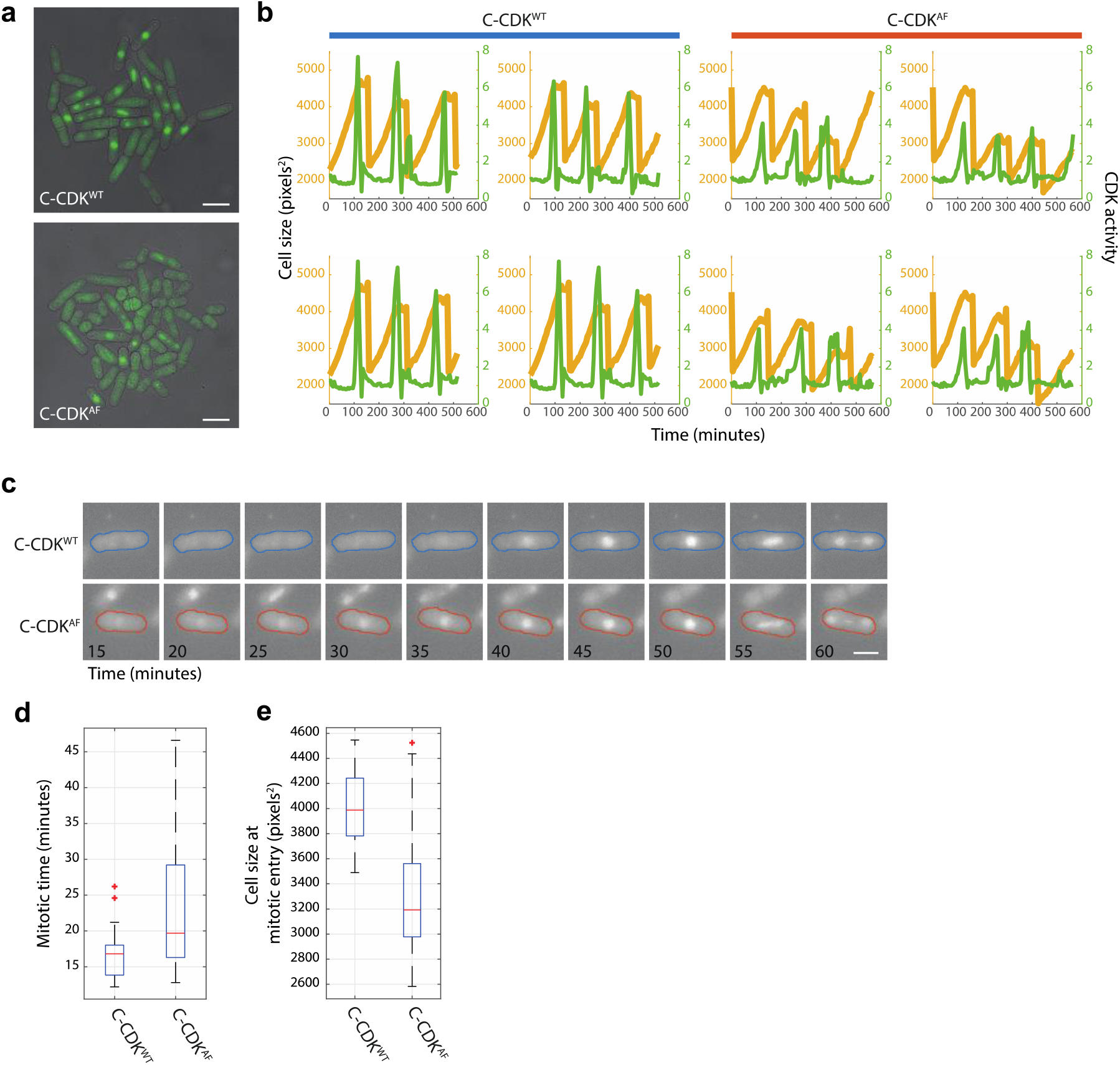
Cut3-GFP as a marker of CDK activity in WT and AF cell strains. **a** Still images of Cut3-GFP tagged in strains expressing C-CDK^WT^ and C-CDK^AF^. Cells were grown in a Cellasics microfluidics device in YE4S at 32°C. Scale bar=10 μm. **b** Example cell length and Cut3-GFP single cell lineages. Quantification is performed by Pomseg and Pomtrack (see methods). Cut3-GFP nuclear/cytoplasmic (N/C) ratio is calculated by dividing mean cytoplasmic Cut3 intensity by mean nuclear Cut3 intensity after background subtraction. Orange lines= cell size, green lines= CDK activity (measured by Cut3 N/C ratio). **c** Montage of tagged C-CDK^WT^ and C-CDK^AF^ strains from time-lapse. Colour outline indicates strain and is derived from Pomseg based segmentation of the brightfield image. Scale bar=5 μm. **d** Boxplot of mitotic times in C-CDK^WT^ and C-CDK^AF^ strains. Mitotic time is calculated as peak time – mitotic entry time. Difference is significant by two sample t-test (p=0.006). Box represents median value delimited by 25^th^ and 75^th^ percentiles. See methods for outlier points. **e** Boxplot of cell size at mitotic entry (cell size sampled at red x position in **Fig. 1i**). Note high variability in the C-CDK^AF^ population (CoV=0.18 vs 0.08 in WT). Box represents median value delimited by 25^th^ and 75^th^ percentiles. See methods for outlier points.

**Supplementary Figure 4:**
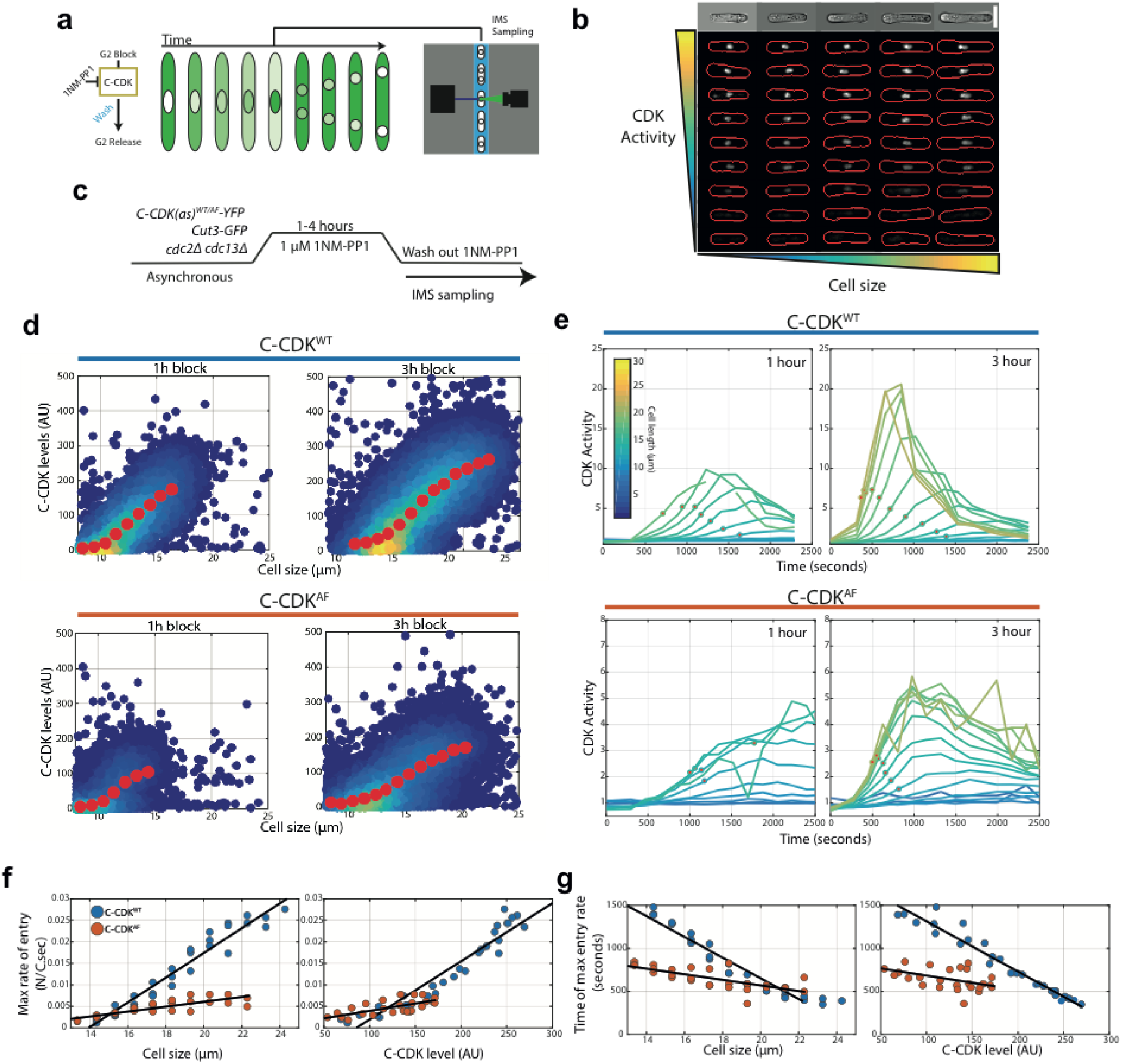
An imaging flow cytometry assay reveals that size, C-CDK level and tyrosine phosphorylation dictate the rate and timing of CDK activation at mitosis. **a** Schematic of the high-throughput imaging flow cytometry block and release assay. Cells are arrested in G2 using 1NM-PP1 for various lengths of time, before being washed of 1NM-PP1 and sampled on an imaging flow cytometer. **b** Representative images of single cells with computed cell masks overlaid on fluorescent Cut3 images in red. Top row of images is from the brightfield channel of the top row of fluorescent images. Representative images taken from Cut3-GFP cells in EMM at 32°C. Scale bar = 10 μm. **c** Experimental outline for panels (D-G). C-CDK^WT/AF^ cells sensitive to the CDK inhibitor 1NM-PP1 are blocked for variable amounts of time. Cells are then washed of 1NM-PP1 and released into mitosis. After release, cells are monitored via sequential sampling using imaging flow cytometry. Block performed using 1 μM 1NM-PP1. Cells were grown in EMM at 32°C. **d** Quantification of C-CDK-YFP levels after indicated block time. Colours indicate density of data; yellow represents high density. Red data points indicate mean of binned data, bin widths 0.33 μm. **e** Plots of mean CDK activity (as measured by Cut3 N/C ratio) within size bins indicated by line colours. Red dots indicate points of maximum Cut3 N/C ratio change, as derived from the first derivative of a smoothing spline fit to raw data (raw data is shown). Each point on line has >50 cells. N=3000-12000 per time point, with ∼400,000 single cell images analysed in total. Background subtraction for N/C ratio performed using wild-type cells lacking Cut3-GFP after indicated block time. **f** Maximum Cut3 N/C ratio change against cell size or C-CDK level. C-CDK level is predicted from data in **d**. Data is taken from 2,3 and 4 hour releases. Black line represents linear regression line. **g** Time of maximum Cut3 N/C ratio change against cell size or C-CDK level. C-CDK level is predicted from data in **d**. Data is taken from 2,3 and 4 hour releases. Black line is the linear regression line. Colours represent the same as panel (F).

**Supplementary Figure 5:**
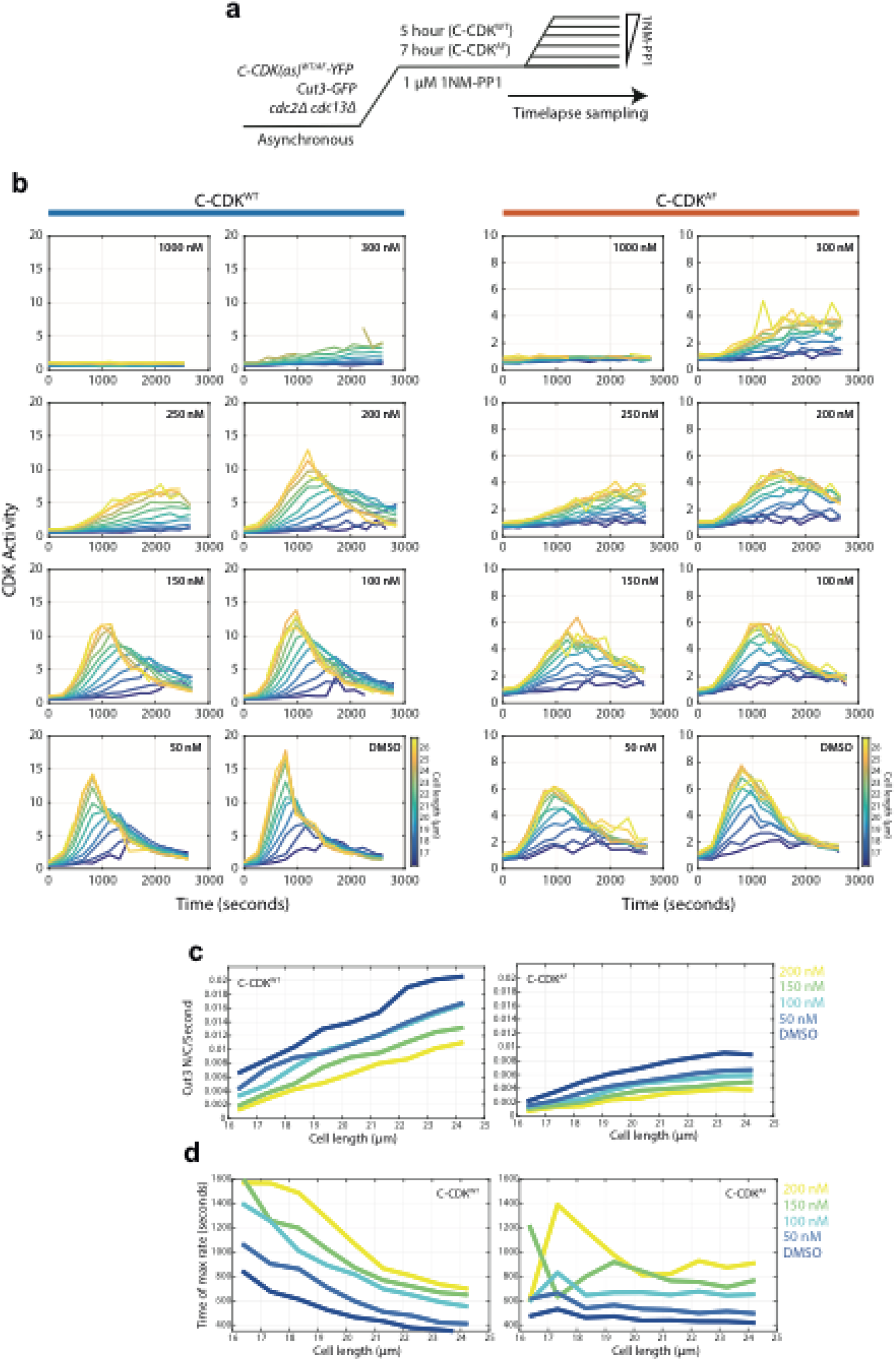
Size dependent grading of mitotic entry rates and timing are dose responsively dependent on CDK inhibition. **a** Experimental outline for panels B-D. 1NM-PP1 sensitive C-CDK^WT^ and C-CDK^AF^ cells are blocked by addition of 1NM-PP1. C-CDK^AF^ cells were blocked for longer (7 hours against 5 hours) to allow cells to reach a similar size distribution to C-CDK^WT^ cells. Cells were then released into a range of 1NM-PP1 concentrations. After release, images were acquired every minute. Time between washing and image acquisition is ∼5 minutes. Cells were grown in EMM at 32°C. Cells are sampled during the region marked time-lapse. **b** Plots of mean CDK activity (as measured by Cut3-GFP N/C ratio) against time from release in indicated size bins at annotated 1NM-PP1 levels. N=1000-4000 cells per time-point, >10 cells averaged within each bin. **c** Plots of maximum Cut3 nuclear translocation rates against cell size in C-CDK^WT^ and C-CDK^AF^ cells. Maximum rates were taken from the first derivative of a smoothing spline fit to data in **b**. Line colours indicate 1NM-PP1 concentration. Key given on the right hand side. **d** Plots of time of maximum Cut3 translocation rate timing vs cell size in WT and AF cells. Maximum rates were taken from the first derivative of a smoothing spline fit to data in **b**. Line colours indicate 1NM-PP1 concentration.

**Supplementary Figure 6:**
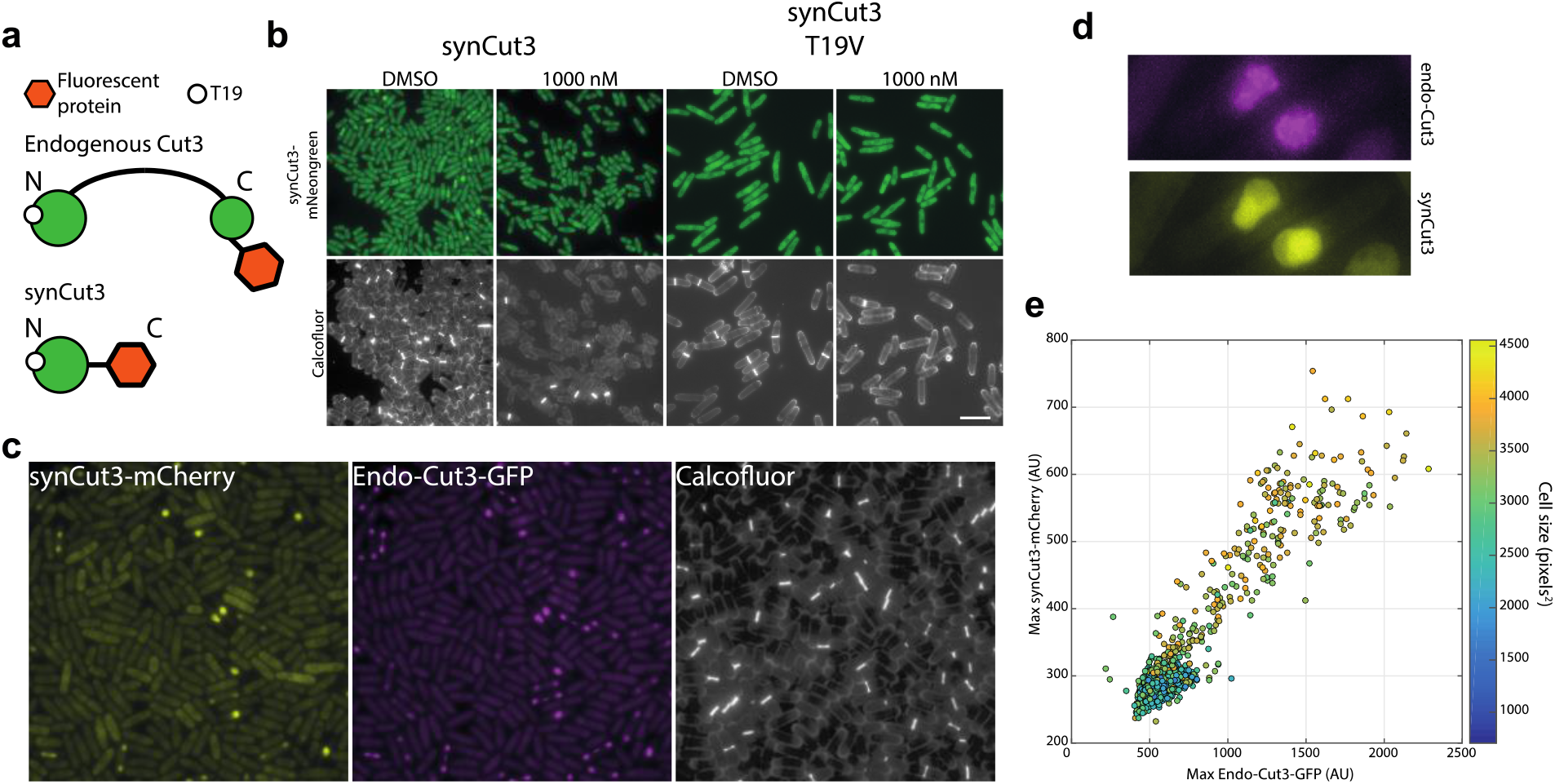
A new synthetic CDK sensor for S. pombe. **a** Design of the synthetic Cut3 (synCut3) sensor. The design includes the first 528 amino acids of Cut3 (and has previously been shown to translocate into the nucleus at mitosis^1^). **b** Example images of synCut3-mNeonGreen expressed from the eno101 promoter, in the presence or absence of 1NM-PP1 (for 1 hour) or a mutated T19 residue. The T19V mutation does not allow CDK phosphorylation, therefore preventing nuclear translocation. Scale bar = 20 μm. **c** Examples images of exogenous synCut3-mCherry and endogenous Cut3-GFP expressing cells. Scale bar = 20 μm. **d** Detailed view of two mitotic cells expressing both synCut3-mCherry and Cut3-GFP. **e** Quantification of exogenous synCut3 signal vs endogenous Cut3 nuclear levels. Data points coloured to indicate cell size. Note endogenous Cut3 signal is smoothed to remove foci containing condensed chromatin regions.

**Supplementary Figure 7:**
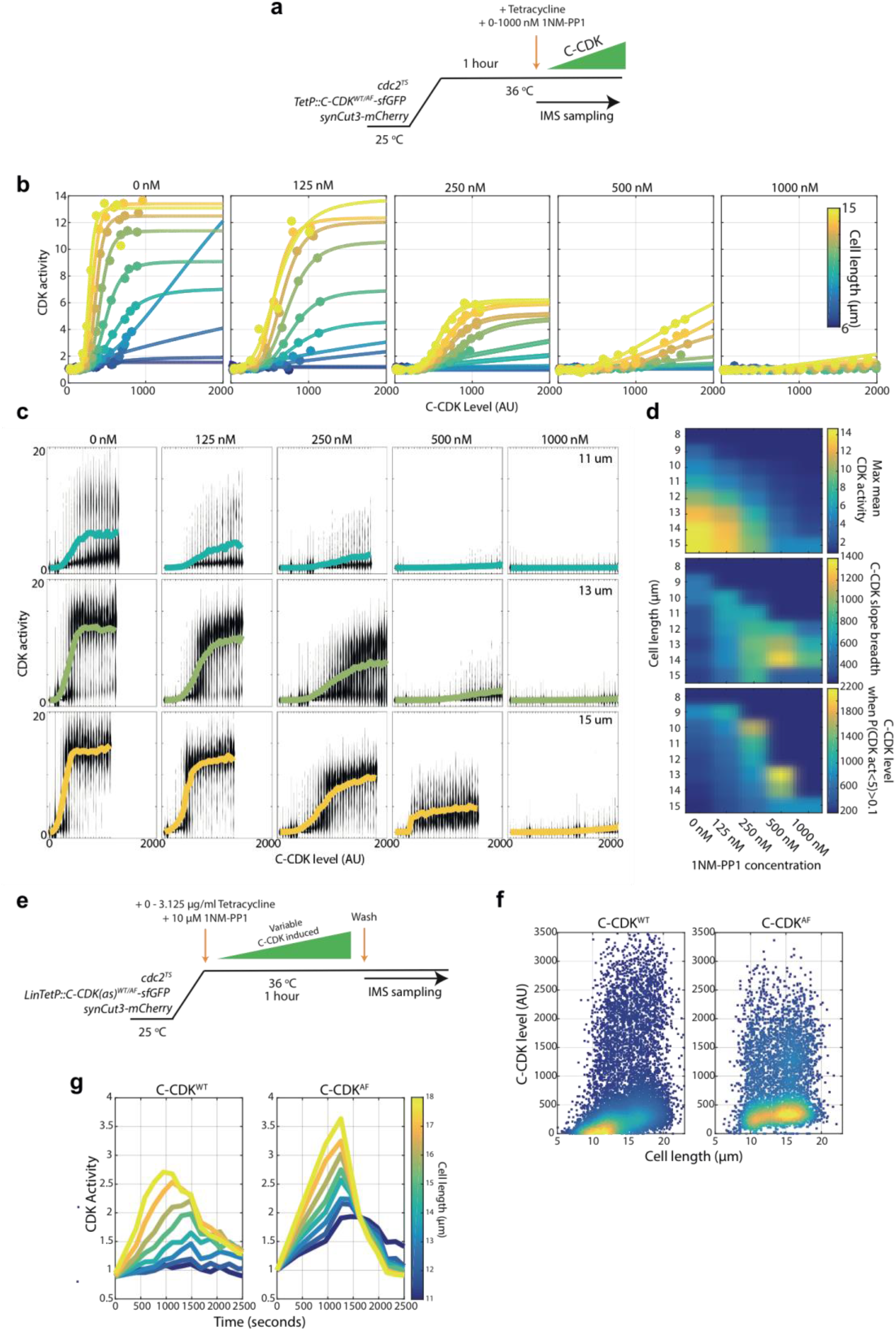
A single cell *in vivo* biochemistry approach permits decoupling of cell size from C-CDK concentration. **a** Experimental outline for panels B-D. Cells were held at 36°C for 1 hour to ablate *cdc2-M26* function. After 1 hour, C-CDK^WT^ or C-CDK^AF^ was induced with tetracycline. Induced C-CDK lacks its degron box sequence, and therefore is not degraded at anaphase. Sequential sampling during C-CDK induction begins at the point of tetracycline addition. Concurrent with tetracycline addition, 1NM-PP1 was added to the specified concentration to inhibit the induced C-CDK. **b** Mean CDK activity against C-CDK level, within specified size bins. Colours within subplot indicate cell size bin (see colour bar). Different subplots represent cells released into different 1NM-PP1 concentrations. **c** Violin plots of single cell C-CDK level against CDK activity data. Individual subplots are the single cell data from a given size bin and 1NM-PP1 level. Rows correspond to the same size bin, columns to the same 1NM-PP1 level. Although bistable behaviour is observed, lines through data represent the population mean C-CDK activity level within a given C-CDK level bin. **d** Heatmap of annotated features, extracted from the single cell dose response data. Max mean CDK activity is the maximum mean CDK activity within a C-CDK fluorescence level bin. C-CDK slope breadth is the change in C-CDK between the C-CDK bin at which CDK activity is greater than 1.1x of minimum, and less than 0.8x of maximum. C-CDK level when P(CDK>5)>0.1 indicates the C-CDK level required to increase CDK activity in 10% of cells to a level greater than 5. **e** Experimental outline for panels F and G. Cells were held at 36°C for 1 hour to ablate *cdc2-M26* function. After 1 hour, C-CDK^WT^ or C-CDK^AF^ was induced with tetracycline to different levels by adding variable amounts of tetracycline. C-CDK was induced in the presence of 10 μM 1NM-PP1 to inhibit the induced C-CDK. After 60 minutes, 1NM-PP1 was washed from cells and cells were sequentially sampled using imaging flow cytometry (IMS). All time measurements are given as time from washing 1NM-PP1. **f** Scatter plot of C-CDK levels against cell size after C-CDK induction. Data represent pooled data from all cells encompassing all 1NM-PP1 release concentrations Colours indicate local data point density. N>10000. **g** synCut3 N/C ratio (representing CDK activity) against time in the presence of induced C-CDK^WT^ or C-CDK^AF^. Line colours indicate size bins. N>50 cells per data point.

